# Transcript-resolved analysis identifies lncRNA carrier architectures and selective nucleolar enrichment in colorectal cancer

**DOI:** 10.64898/2026.11.09.765366

**Authors:** Bridget Flowers, Penelope Lialios, Eileen Park, Chelsea Denniss, Rebecca J. Burgess, Jong-Sun Lee

**Affiliations:** Center for Genetic Diseases, Chicago Medical School, Rosalind Franklin University of Medicine and Science, North Chicago, IL 60064, USA; Division of Biochemistry and Molecular Biology, Department of Biomedical Sciences, Southern Illinois University School of Medicine, Carbondale, IL 62901, USA; Department of Biochemistry and Molecular Biology, Chicago Medical School, Rosalind Franklin University of Medicine and Science, North Chicago, IL 60064, USA

## Abstract

Long noncoding RNAs (lncRNAs) in colorectal cancer (CRC) are commonly characterized at the gene level, obscuring processed RNA architectures and regional localization. In renal cell carcinoma, a processed *CRNDE* isoform retaining an ultraconserved element (UCE) with a C/D snoRNA-like region promotes pre-60S maturation. Here, we examined *CRNDE* UCE-region representation and integrated CRC tumor expression and clinical evidence with published HEK293 cross-linking and immunoprecipitation (CLIP) profiles for C/D small nucleolar ribonucleoprotein (snoRNP) proteins and long-read sequencing to identify retained CLIP-supported regions. *CRNDE* was upregulated in 49 of 50 paired tumors, and UCE-region signal was detected in 380 of 382 assessable primary tumors, although total *CRNDE* abundance showed little association with stage or outcome. Processed UCE-retaining *CRNDE* transcripts were identified in Caco-2 cells. Integration identified 1,714 disease-supported lncRNAs with CLIP overlap; among 576 non-*CRNDE* candidates, 188 CLIP-supported regions were retained within processed transcripts from 65 loci in both CRC cell models. Region-directed RT-qPCR following pre-ribosome sequential extraction revealed preferential enrichment of selected *JPX*, *DLEU2*, and *LINC00630* RNA populations in the core nucleolar fraction relative to broader transcript-region measurements in both cell lines. These findings identify retained-region transcript architectures and reveal region-dependent nucleolar enrichment not apparent from broader transcript measurements.

## 1. Introduction

Colorectal cancer (CRC) is among the most commonly diagnosed malignancies worldwide and remains a leading cause of cancer-related mortality^1^. Sustained tumor growth and proliferation impose increased demands on protein synthesis and, consequently, on the production of functional ribosomes^2^. Ribosome biogenesis is a highly coordinated process that begins in the nucleolus, where ribosomal RNA (rRNA) transcription, processing, modification, and assembly with ribosomal proteins initiate the formation of ribosomal subunits^2^. Oncogenic pathways, including MYC and mTOR signaling, stimulate ribosome production to support the translational demands of malignant cells^3,4^. In CRC, dysregulation of ribosome biogenesis has been linked to oncogenic signaling and altered translational programs, and experimental inhibition of rRNA transcription can suppress tumor cell proliferation and promote differentiation^2,5^.

Noncoding RNAs play essential roles in ribosome biogenesis, particularly small nucleolar RNAs (snoRNAs), which direct site-specific modification and processing of precursor rRNA (pre-rRNA)^6,7^. SnoRNAs are broadly classified into two major families: box C/D snoRNAs, which guide 2′-O-methylation, and box H/ACA snoRNAs, which direct pseudouridylation^6^. Box C/D snoRNAs associate with a conserved protein complex comprising fibrillarin (FBL), NOP56, NOP58, and SNU13 to form functional small nucleolar ribonucleoprotein complexes (snoRNPs)^7^. Beyond their roles in rRNA modification, specific snoRNAs also participate directly in pre-rRNA cleavage and maturation, coordinating multiple steps of ribosome biogenesis^7^.

In mammals, many snoRNAs are encoded within introns of protein-coding or long noncoding RNA (lncRNA) genes and are released through splicing-dependent processing followed by exonucleolytic trimming^7^. For example, the lncRNA *GAS5*, which contains intronic box C/D snoRNAs, is a well-characterized example of snoRNA-host processing^8^. SnoRNA-related sequences also occur in lncRNAs that regulate nucleolar processes. *SLERT*, a snoRNA-ended lncRNA, promotes pre-rRNA transcription by modulating the RNA helicase DDX21, whereas LoNA contains a snoRNA-like region that interacts with FBL and negatively regulates rRNA methylation^9,10^. More recently, an alternatively spliced isoform of *CRNDE* containing an ultraconserved element (UCE) with a C/D snoRNA-like region, termed *CRNDE*^UCE^, was shown to promote 60S ribosomal subunit biogenesis in renal cell carcinoma (RCC)^11^. Unlike conventional intronic snoRNAs that are released from their host transcript, the C/D snoRNA-like element of *CRNDE*^UCE^ remains embedded within a longer processed transcript, where it interacts with pre-rRNA and facilitates recruitment of the 60S biogenesis factor eIF6^11^. This configuration exemplifies a retained-region RNA carrier architecture, in which a defined RNA element remains embedded within a longer processed transcript rather than being excised.

Although *CRNDE* was originally identified through its marked upregulation in colorectal neoplasia, whether its UCE-containing functional region is retained within processed transcripts in CRC remains unclear^12^. Neither total *CRNDE* expression nor short-read coverage of the UCE-containing region can establish which processed RNA molecules retain this region. Moreover, whether other CRC-associated lncRNAs retain regions overlapping published C/D snoRNP-protein CLIP signals within processed transcripts remains largely unexplored. Here, guided by the functional precedent of *CRNDE*^UCE^, we characterized *CRNDE* expression, UCE-region representation, and processed transcript architecture in CRC. We then integrated primary tumor expression and clinical associations with published CLIP profiles of C/D snoRNP proteins generated in HEK293 cells and long-read RNA sequencing to identify additional CRC-associated lncRNAs retaining CLIP-supported regions within processed transcripts. Finally, we examined the subcellular distribution of selected RNA populations using pre-ribosome sequential extraction (PSE) and region-directed RT-qPCR. Together, these analyses identify a broader landscape of CRC-associated lncRNAs retaining CLIP-supported regions within processed transcripts and reveal preferential core nucleolar enrichment of selected RNA populations that is not apparent from broader transcript-region measurements of the same loci.

## 2. Results

### 2.1. CRNDE is recurrently upregulated in colorectal cancer, and its UCE-containing region is widely detected in primary tumors

*CRNDE* was originally identified through its marked upregulation in colorectal neoplasia^12^, a finding subsequently reproduced in independent CRC cohorts and supported by functional studies implicating *CRNDE* in tumor growth and other oncogenic phenotypes^13,14^. To define its patient-level expression pattern, we analyzed RNA sequencing (RNA-seq) data from The Cancer Genome Atlas (TCGA) COAD and READ cohorts using strictly matched tumor and adjacent normal samples (Table S1). *CRNDE* expression was higher in the tumor tissue in 49 of 50 patients, with an overall log2 fold change of 3.98 (Figure 1A), consistent with its previously reported upregulation in CRC. Previous studies have associated high *CRNDE* expression with advanced disease and adverse clinical outcome in CRC, including reports of increased expression in advanced-stage tumors and poorer overall or disease-free survival^14,15^. In the TCGA COAD and READ cohorts, however, *CRNDE* transcript abundance showed little association with progression-free interval, overall survival or pathologic stage (Figure S1A; Table S2). These results distinguish the strong upregulation of *CRNDE* in CRC from its more limited association with disease progression and clinical outcome.

**Figure 1.**
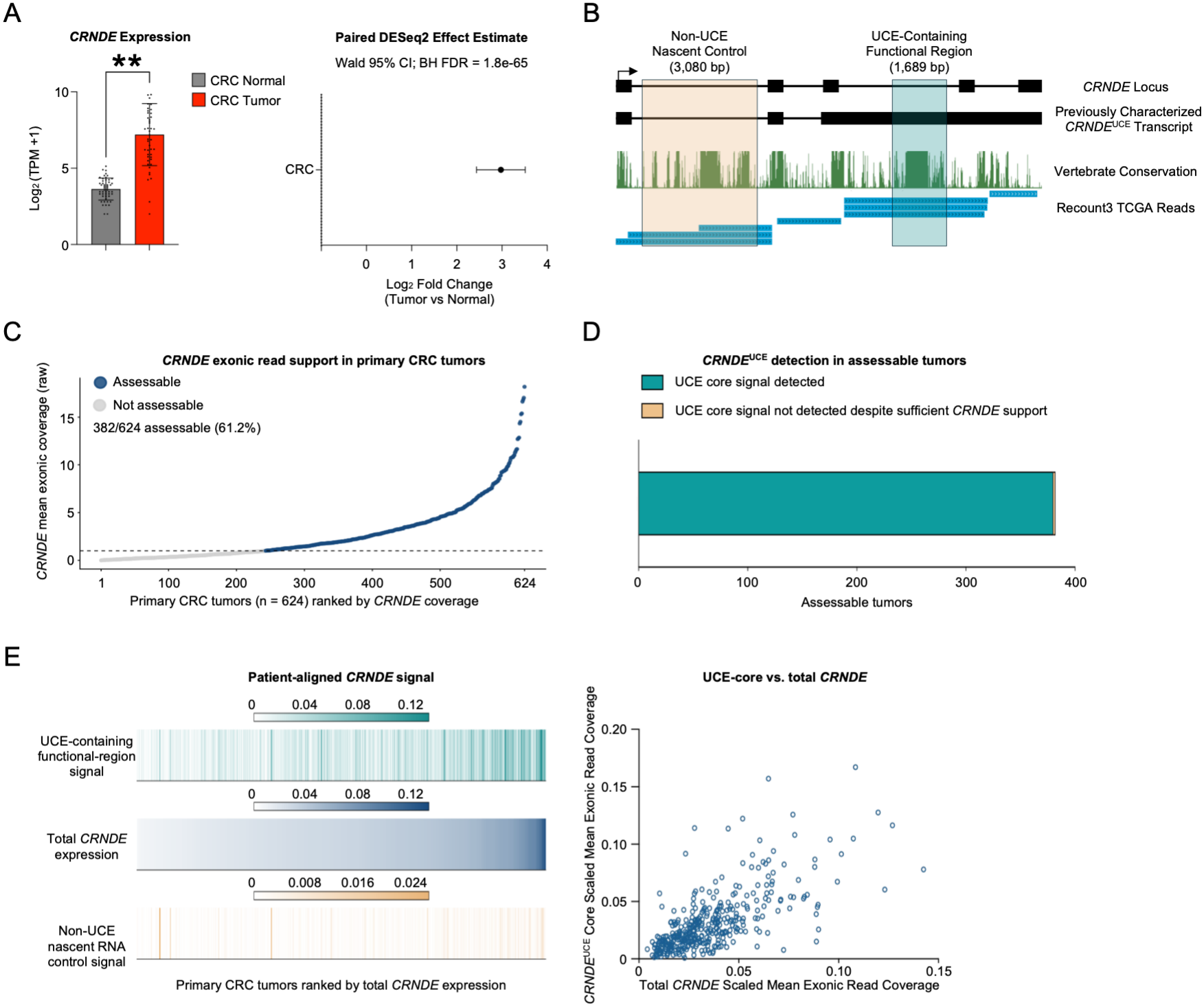
*CRNDE* is recurrently induced in colorectal cancer and the *CRNDE*^UCE^ functional core is broadly detected in primary tumors. (A) Paired tumor-normal differential expression of *CRNDE* in CRC. Bar chart (left) shows *CRNDE* expression in paired CRC tumor and normal samples (n = 50); forest plot (right) shows effect estimate and 95% confidence interval. Differential expression was analyzed using DESeq2 with apeglm-shrunken log2 fold-change estimates. **P <0.01, calculated by two-tailed t-test. (B) *CRNDE* genomic architecture and RNA-seq measurement strategy. UCSC genome browser ENCODE transcription, vertebrate conservation (PhastCons), and RNA sequencing read coverage (recount3) aligned with previously characterized *CRNDE*^UCE^ transcript. (C) *CRNDE* exonic read support in primary CRC tumors. Dot plot shows *CRNDE* mean exonic raw coverage vs. 624 CRC tumor samples ranked by *CRNDE* coverage. Samples were considered *CRNDE*-assessable if raw *CRNDE* mean exonic coverage ≥ 1x. (D) *CRNDE*^UCE^ in assessable primary CRC tumors. The UCE core signal was detected in 380/382 assessable tumors. (E) Patient-aligned *CRNDE* signal. UCE-containing functional-region signal, total *CRNDE* expression, and non-UCE nascent RNA control signal are shown for tumors ranked by total *CRNDE* expression from recount3 (left). Scatterplot of *CRNDE* UCE-region versus total *CRNDE* scaled mean exonic read coverage (right). UCE-region coverage was positively correlated with total *CRNDE* coverage, although relative UCE-region representation varied across tumors.

The recent identification in RCC of an alternatively spliced, approximately 5-kb, three-exon *CRNDE* isoform containing an ultraconserved element (UCE), termed *CRNDE*^UCE^, raised the question of whether *CRNDE* isoforms containing the same RNA region are expressed in CRC^11^. We therefore analyzed recount3-derived short-read RNA-seq coverage from primary CRC tumors across the previously characterized 1,689-nt UCE-containing region, all *CRNDE* exons and a non-UCE intronic control region (Figure 1B and Table S1). Of 624 CRC tumors, 382 had sufficient *CRNDE* exonic read coverage for regional assessment (Figure 1C), and the UCE-containing region was detected in 380 of these tumors (99.5%; Figure 1D). Adjacent normal tissue generally lacked sufficient *CRNDE* read support for regional assessment. Among assessable tumors, UCE-region coverage was positively correlated with total *CRNDE* exonic coverage (Spearman *ρ* = 0.69, *P* = 2.27 x 10^-55^), although the relative representation of the UCE-containing region varied across tumors (Figure 1E; Table S1). Thus, RNA corresponding to the UCE-containing region was detected in nearly all assessable primary CRC tumors, although short-read RNA-seq could not establish whether this region was retained within processed *CRNDE* transcripts.

We next examined whether variation in UCE-region signal was associated with transcriptional programs consistent with the ribosome-biogenesis function previously established for *CRNDE*^UCE^ in RCC^11^. Gene-expression changes associated with UCE-region signal showed a positive directional association with ribosome-biogenesis genes, but the prespecified gene set did not reach statistical significance, and the association was not distinguishable from that observed for matched intronic control regions (Figure S1B and Table S3). Thus, the primary tumor data did not establish a UCE-specific ribosome-biogenesis transcriptional program in CRC.

### 2.2. Long-read RNA-seq identifies processed CRNDE transcripts retaining the UCE-containing region and reveals nucleolar enrichment of UCE-region RNA in CRC cells

Although short-read RNA-seq detected the *CRNDE* UCE-containing region in nearly all assessable primary CRC tumors, it could not establish whether this region was retained within processed *CRNDE* transcripts. We therefore examined individual *CRNDE* molecules using publicly available long-read RNA-seq datasets. We defined a processed *CRNDE*^UCE^ carrier as a primary, same-strand *CRNDE* alignment with a mapping quality ≥20, retention of ≥95% of the 1,689-nt UCE-containing region, and at least one independent splice junction outside that interval (Figure 2A). These criteria permitted identification of both RCC-like^11^ and alternative processed *CRNDE* transcripts retaining the same functional region.

**Figure 2.**
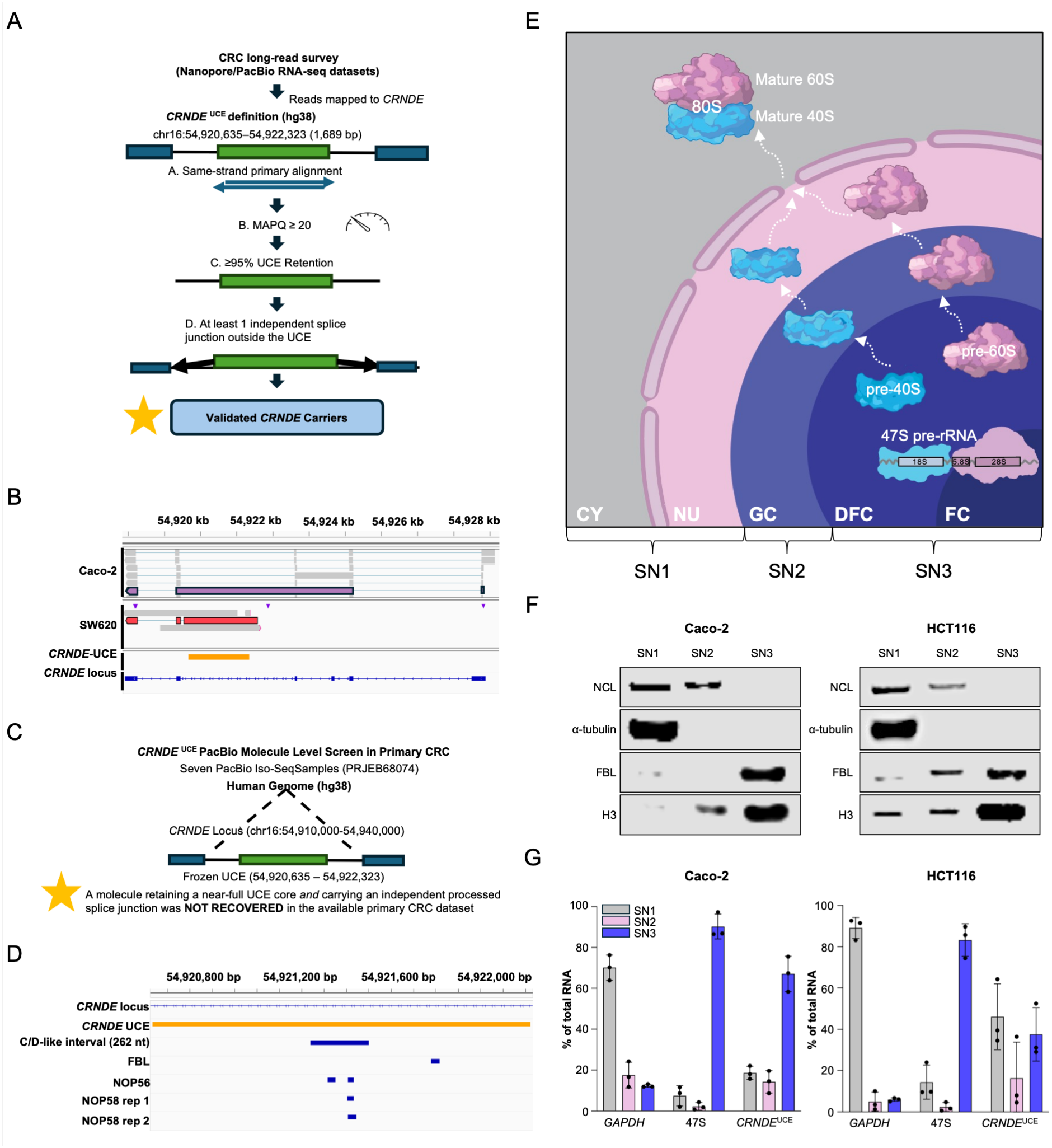
Long-read RNA-seq analysis identifies processed *CRNDE*^UCE^ carriers in CRC with orthogonal support for C/D snoRNP association. (A) Schematic showing filtering from CRC long-read survey of Nanopore/PacBio datasets to validated *CRNDE* carriers. Publicly available Nanopore and PacBio long-read RNA-sequencing datasets were screened for transcripts mapping to the CRNDE locus. Processed *CRNDE* UCE carriers were identified using four criteria: (i) primary alignment concordant with the *CRNDE* transcriptional strand, (ii) mapping quality (MAPQ) ≥20, (iii) retention of ≥95% of the UCE-containing region and (iv) at least one independent splice junction outside the UCE interval. These criteria distinguish processed UCE-retaining transcripts from unspliced or UCE-excising *CRNDE* architectures. (B) Representative genome-browser tracks illustrate the recovered transcript architectures relative to the *CRNDE* locus and UCE-containing interval (GRCh38), demonstrating recovery of processed UCE-retaining *CRNDE* transcripts in CRC cell-line models. (C) Schematic showing the *CRNDE*^UCE^ PacBio molecule level screen in primary CRC. Seven publicly available PacBio Iso-Seq specimens comprising tumor and adjacent normal tissues (PRJEB68074) were analyzed for *CRNDE* transcript architectures using the predefined UCE-retention and independent splice-junction criteria. Among 21 high-confidence, same-strand *CRNDE* molecules, 12 exhibited splice junctions excising the UCE-containing interval, three partially overlapped the UCE without independent splice support, and six lacked detectable UCE overlap without meeting the excision criterion. No molecules retained ≥95% of the UCE-containing region, and no qualifying processed *CRNDE* UCE carrier was recovered at the available sequencing depth. (D) IGV snapshot of the *CRNDE* locus showing CLIP overlap of FBL, NOP56, and NOP58. Two NOP56 peaks overlapped the C/D-like interval by 34 and 27 nt, respectively, while NOP58 peaks from two independent libraries overlapped the interval by 29 and 38 nt. FBL did not directly overlap the C/D-like interval but exhibited a 37-nt peak elsewhere within the broader UCE-containing region. These observations provide independent CLIP-based support for NOP56 and NOP58 association with the previously characterized *CRNDE* C/D-like element. (E) Schematic illustration of sequential pre-ribosomal extraction highlighting the stages of ribosome biogenesis represented by each fraction (SN3, SN2, and SN1). Cy, cytoplasm; Nu, nucleoplasm; GC, granular component; DFC, dense fibrillar component; FC, fibrillar center. (F) Western blot analysis of representative immunoblots of nucleolin (NCL), α-tubulin, fibrillarin (FBL), and histone H3 across sequential extraction fractions from Caco-2 and HCT116 cells: SN1, SN2, and SN3. (G) Caco-2 and HCT116 cells RT-qPCR analysis of *CRNDE*^UCE^ in nucleolar fractions. *GAPDH* and 47S pre-rRNA represent cytoplasmic and early nucleolar markers, respectively. Data are presented as mean ± SD (n=3 biological replicates).

Among the seven CRC cell lines investigated, Caco-2 provided the strongest evidence of processed *CRNDE*^UCE^ carrier transcripts. Of 249 primary same-strand *CRNDE* reads, 246 were assessable for UCE retention and processing status. Eight retained ≥95% of the UCE-containing region, of which four also contained an independent splice junction outside the UCE, meeting the processed-carrier definition (Figure 2B). The remaining four retained the UCE-containing region but lacked an independent splice junction, distinguishing regional retention from evidence of a processed carrier. All four qualifying carriers were recovered from one of two independent Caco-2 datasets (Figure S2A). SW620 cells yielded one additional qualifying molecule among three high-confidence *CRNDE* reads, whereas no qualifying carriers were recovered from HCT116, LoVo, or SW480, which yielded 32, five, and two high-confidence reads, respectively (Figure S2B). Colo320 and LS174T lacked high-confidence read support and were unassessable. At the observed Caco-2 carrier frequency of 4/246 (1.6%), approximately 100 assessable *CRNDE* reads would be required for an 80% probability of recovering at least one carrier, assuming independent sampling and a comparable carrier frequency. Of the CRC cell lines with available long-read expression data, only Caco-2 reached this approximate sampling depth. Thus, processed UCE-retaining *CRNDE* transcripts were identified in Caco-2 cells, with an additional qualifying molecule detected in SW620, whereas limited molecular sampling precluded reliable assessment of carrier prevalence across the other CRC models.

We next applied the same molecule-level criteria to PacBio Iso-Seq data from seven primary CRC-associated specimens, comprising tumor and adjacent normal tissue (Figure 2C). Following Iso-Seq-specific strand-orientation handling, 21 high-confidence, same-strand *CRNDE* molecules were identified, 19 of which originated from a single tumor specimen. Of these 21 molecules, 12 contained splice junctions removing the UCE-containing interval, three partially overlapped the UCE without an independent splice junction, and six lacked detectable UCE overlap without meeting the excision criterion. None retained ≥95% of the UCE-containing region. Thus, no qualifying processed *CRNDE*^UCE^ carrier was recovered from these primary CRC-associated specimens, but the limited molecular sampling precluded reliable estimation of carrier prevalence in primary CRC tumors.

The association of *CRNDE*^UCE^ with C/D snoRNP components was previously established in RCC^11^. To examine independent CLIP evidence for the same RNA region, we analyzed published transcriptome-wide CLIP datasets generated in HEK293 cells for the C/D snoRNP proteins NOP56, NOP58, and FBL^16^, focusing on the previously defined *CRNDE*^UCE^ C/D-like interval (Figure 2D, Table S4). Two NOP56 CLIP peaks overlapped this interval by 34 and 27 nt, respectively. NOP58 provided replicate-consistent support, with peaks from two independent libraries overlapping the same interval by 29 and 38 nt. FBL did not directly overlap the C/D-like interval, although a 37-nt FBL peak was detected elsewhere within the broader UCE-containing region. These independent CLIP data provide additional evidence consistent with NOP56 and NOP58 association with the *CRNDE*^UCE^ C/D-like region, although they do not directly establish binding in CRC cells. AlphaFold3^17^ modeling further suggested structural compatibility with a C/D snoRNP-like assembly (Figure S2C), without establishing native complex formation or experimentally validated protein-RNA interactions.

Having identified processed UCE-retaining *CRNDE* transcripts in CRC cells, we next examined the subcellular distribution of RNA containing the UCE region. We performed pre-ribosome sequential extraction (PSE) in Caco-2 and HCT116 cells to separate cyto-nuclear components (SN1), outer nucleolar components associated with later stages of pre-ribosome maturation (SN2), and core nucleolar components enriched for early pre-ribosomal species (SN3)^18,19^ (Figure 2E). Western blot analysis demonstrated the expected distribution of compartment-specific protein markers (Figure 2F), while RT-qPCR confirmed preferential recovery of *GAPDH* in SN1 and 47S pre-rRNA in SN3 (Figure 2G), supporting successful fractionation. RT-qPCR targeting the *CRNDE* UCE-containing region revealed preferential enrichment of UCE-region-containing RNA in SN3 in Caco-2 cells, whereas the corresponding signal was distributed between SN1 and SN3 in HCT116 (Figure 2G). Together, these findings establish the presence of processed *CRNDE* transcripts retaining the UCE-containing region in CRC cells and reveal preferential core nucleolar enrichment of UCE-region-containing RNA in Caco-2 cells. These observations extend the molecular characterization of the *CRNDE* UCE region beyond RCC and provide a basis for investigating whether related RNA-region retention features occur in other CRC-associated lncRNAs.

### 2.3. Integration of snoRNP-protein CLIP overlap with CRC disease evidence identifies candidate lncRNAs

The identification of a C/D snoRNA-like region retained within processed *CRNDE*^UCE^ transcripts prompted us to investigate whether other lncRNAs contain regions overlapping C/D snoRNP-protein CLIP signals. We therefore mapped published FBL, NOP56, and NOP58 CLIP data^16^ generated in HEK293 cells across 34,866 GENCODE-annotated lncRNAs using same-strand exonic overlap (Figure 3A). Of these, 9,447 lncRNAs overlapped at least one snoRNP-protein CLIP signal, including 4,067 with overlap from at least two proteins and 1,764 with overlap from all three (Figure 3A; Table S4). Increasing the minimum overlap requirement to ≥10 nt in a sensitivity analysis retained 9,285 of these lncRNAs (Figure S3A). However, overlap with multiple proteins did not exceed expectations under a degree-preserving null model accounting for overall CLIP exposure, whereas overlap in both independent NOP58 libraries showed a modest but significant enrichment (Figure S3B).

**Figure 3.**
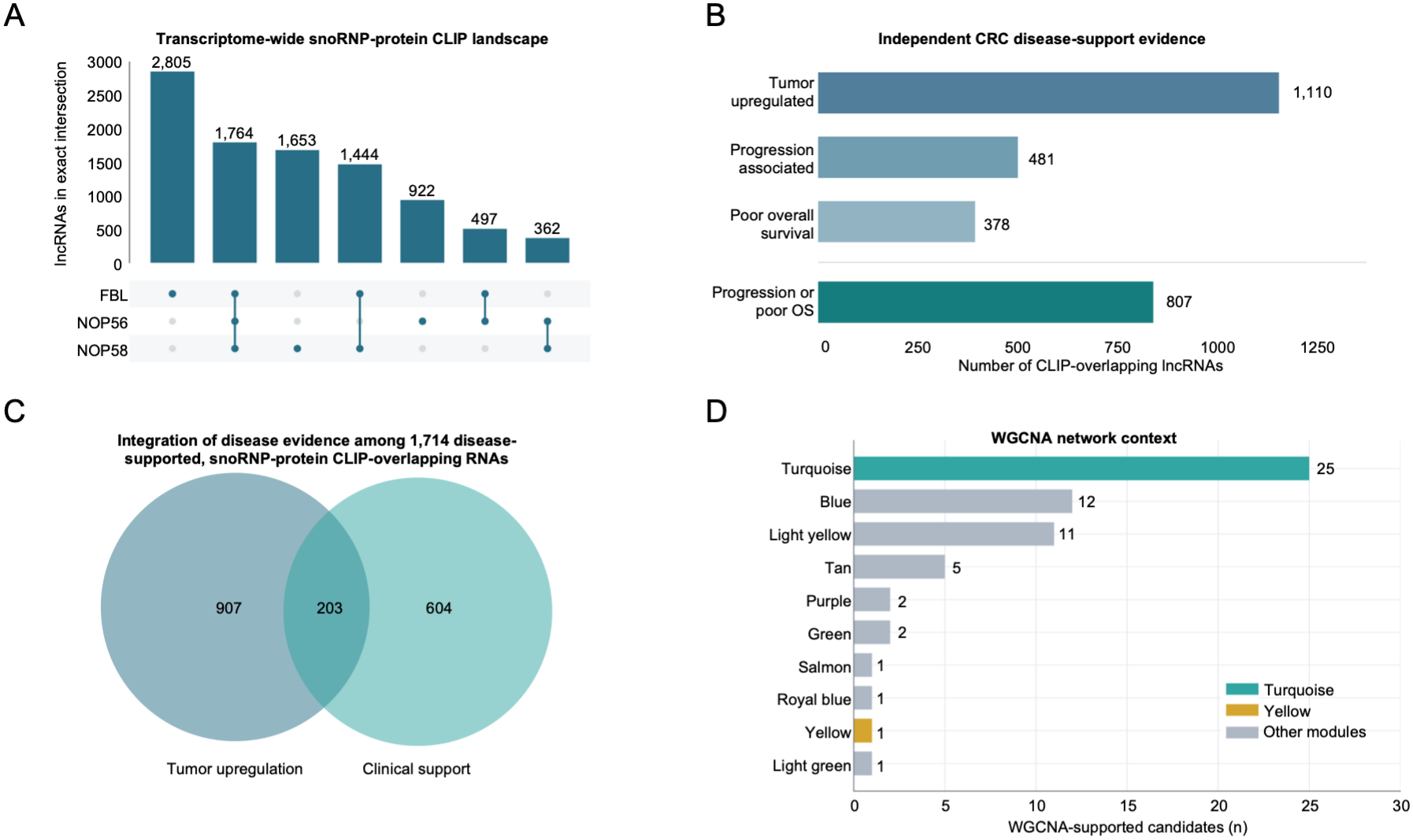
Integration of snoRNP-protein CLIP overlap with independent CRC disease evidence identifies candidate lncRNAs. (A) UpSet plot of transcriptome-wide snoRNP-protein CLIP landscape shows lncRNAs with CLIP overlap for FBL, NOP56, and NOP58. CLIP support was defined by same-strand exonic overlap of ≥1 nucleotide, with either NOP58 replicate considered sufficient for protein-level overlap. Bars indicate the numbers of lncRNAs within each mutually exclusive protein-overlap combination, and the connected-dot matrix identifies the corresponding proteins. Overall, 9,447 lncRNAs overlapped at least one core snoRNP protein, including 4,067 with overlap from at least two proteins and 1,764 with overlap from all three. Overlap counts represent CLIP-supported genomic associations rather than experimentally validated snoRNP binding or function. (B) Independent CRC disease-support evidence (tumor upregulation, progression association, poor overall survival, and progression association or poor overall survival) among CLIP-overlapping lncRNAs. Horizontal bar plot summarizing independent CRC disease evidence among the 9,447 lncRNAs with at least one snoRNP-protein CLIP overlap. Of these, 1,110 were significantly upregulated in CRC tumors relative to normal tissue, 481 were associated with disease progression, and 378 were associated with poor overall survival. The union of progression-associated and poor-survival-associated lncRNAs comprised 807 unique genes with at least one adverse clinical association. Categories are not mutually exclusive, and counts represent gene-level associations rather than distinct transcript isoforms. (C) Integration of disease evidence among 1,714 disease-supported, snoRNP-protein CLIP-overlapping lncRNAs shows 907 tumor-upregulated candidates, 604 clinical candidates with clinical support, and 203 candidates with both clinical support and tumor upregulation. (D) WGCNA network context of candidate lncRNAs. Horizontal bar plot showing the distribution of 61 WGCNA-supported lncRNAs among the final 1,714 CRC disease-supported candidates across ten co-expression modules. Bars represent the numbers of supported candidates assigned to each module. The turquoise and yellow modules are highlighted, containing 25 and one supported lncRNA, respectively; the remaining modules are shown in grey. These modules provide complementary transcriptional-network context and were not used to define membership in the disease-supported candidate population.

We next examined whether these CLIP-overlapping lncRNAs were associated with CRC using gene expression and clinical outcome data. Among the 9,447 CLIP-overlapping lncRNAs, 1,110 were significantly upregulated in CRC tumors relative to matched normal tissue, 481 were associated with disease progression, and 378 were associated with poor overall survival; the latter two analyses identified 807 unique lncRNAs with at least one adverse clinical association (Figure 3B; Table S5). Integration of tumor-associated upregulation with progression or survival association identified 1,714 unique snoRNP-protein CLIP-overlapping lncRNAs with at least one form of independent CRC disease support, comprising 907 with tumor upregulation only, 604 with adverse clinical association only, and 203 with both forms of evidence (Figure 3C; Table S5).

To provide additional transcriptional-network context for the disease-supported lncRNAs, we examined their representation within previously constructed weighted gene co-expression network analysis (WGCNA) modules derived from pooled CRC, gastric cancer, and esophageal cancer tumors^20^. Among the 1,714 candidates, 61 had WGCNA support across 10 modules (Figure 3D; Table S5). Of these, 26 mapped to two modules (25 in the turquoise module and one in the yellow module) enriched for ribonucleoprotein-complex biogenesis, ribosome biogenesis, and rRNA processing. The turquoise module was additionally enriched for cytoplasmic translation and ribosome assembly (Figure S3C). These results placed a subset of the disease-supported lncRNAs within co-expression networks associated with ribosome biogenesis and RNP biology, without using WGCNA module membership as a candidate-selection criterion.

As a supplementary assessment, we tested replicate-consistent NOP58 CLIP overlap among the disease-supported candidates. Replicate-consistent NOP58 overlap was modestly enriched after accounting for overall CLIP exposure (540 observed versus 494.4 expected; empirical *P* = 0.000999; Figure S3D), indicating that the enrichment was not explained solely by differences in overall CLIP exposure. However, this analysis did not account for transcript abundance. Together, these analyses identified 1,714 CRC disease-supported lncRNAs with snoRNP-protein CLIP overlap for subsequent examination of RNA processing and retention within processed transcripts.

### 2.4. CRC cell-line expression and long-read analysis identify processed lncRNAs retaining snoRNP-protein CLIP-supported regions

We next examined the expression of the 1,714 CRC disease-supported lncRNAs across 81 CRC cell lines using transcriptomic data from the Cancer Cell Line Encyclopedia (CCLE)/DepMap ^21,22^. Of the 1,714 candidates, 1,713 were quantified, and 1,359 (79.3%) were expressed at ≥1 TPM in at least one CRC model (Figure 4A; Table S6). To assess their representation relative to other lncRNAs, we compared candidate detection with that of the remaining qualified lncRNAs. Among 15,455 noncandidate lncRNAs, 5,504 (35.6%) met the same expression threshold, corresponding to a 2.23-fold difference in the unadjusted comparison (Figure 4B).

**Figure 4.**
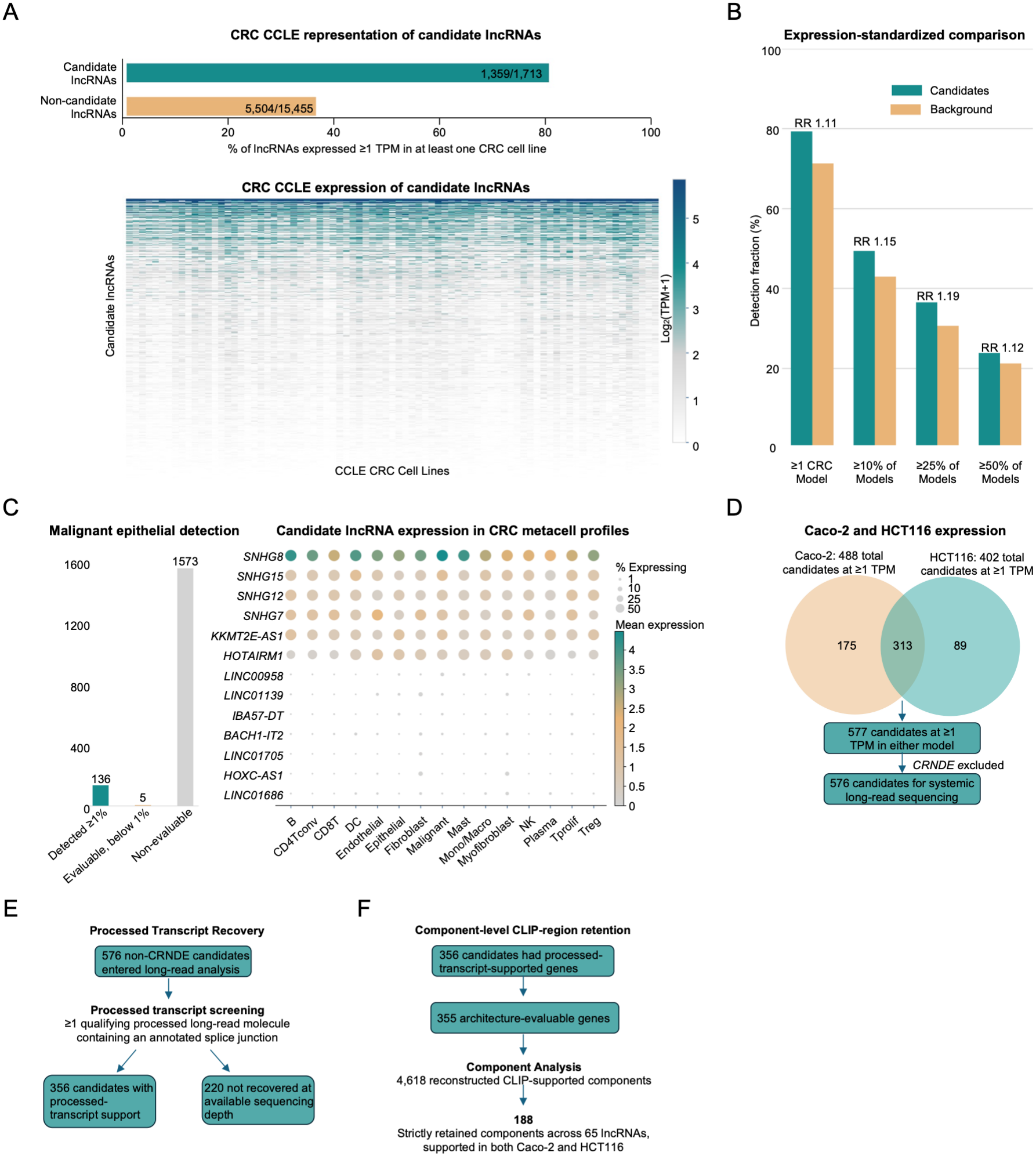
CRC cellular expression and long-read analysis identify transcript-resolved lncRNA carriers. (A) CRC CCLE representation of candidate lncRNAs. Bar chart (top) shows percent of lncRNAs expressed ≥ 1 TPM in at least one CRC cell line. Heat map (bottom) shows distribution of candidate lncRNA expression across 81 CRC CCLE cell lines. (B) Expression-standardized comparison of lncRNA candidates and background in 81 CRC CCLE cell lines. Chart depicts detection fraction of candidates and background lncRNAs and relative risk across ≥ 1 CRC model, ≥ 10% of models, ≥ 25% of models, and ≥ 50% of the 81 CRC CCLE cell line models. (C) Evaluation of candidate lncRNA detection in malignant epithelial single-cell derived metacell profiles from TabulaTIME. Bar chart (left) shows candidate lncRNA detection and evaluation. Dot plot (right) shows selected candidate lncRNA expression across cell types. (D) Caco-2 and HCT116 expression of candidate lncRNAs. Venn diagram and flow chart show selection scheme resulting in 576 candidate lncRNAs proceeding to systematic long-read analysis. (E) Processed transcript recovery of candidates entering long-read analysis. 356 candidates had ≥ 1 qualifying processed long-read molecule containing an annotated splice junction while 220 were not recovered at available sequencing depth. (F) Component-level CLIP-region retention. Of 356 candidates with processed-transcript support, 355 were architecture evaluable, resulting in 4,618 reconstructed CLIP-supported components and 188 strictly retained components across 65 lncRNAs supported in both Caco-2 and HCT116.

Because primary-tumor expression contributed to candidate selection, we next examined whether differences in RNA abundance could account for the higher detection rates of disease-supported candidates in CRC cell lines. We standardized the background lncRNA population to the primary CRC tumor expression distribution of the candidates (Figure S4A). After standardization, 79.3% of candidates and 71.2% of background lncRNAs were expressed in at least one CRC model (RR = 1.11; Figure 4B). Similar results were observed when requiring expression in ≥10%, ≥25%, and ≥50% of the 81 CRC models (RR = 1.15, 1.19, and 1.12, respectively). The number of cell lines expressing individual candidates also varied considerably (Figure S4B). Thus, accounting for primary-tumor expression substantially attenuated the difference observed in the unadjusted comparison, indicating that baseline abundance contributed to the higher detection rates of the disease-supported candidates. This analysis assessed expression-related bias without altering the candidate set.

To determine whether the disease-supported lncRNAs were also detectable in malignant epithelial cells, we analyzed the TabulaTIME single-cell atlas^23^. Among 141 candidates with evaluable malignant epithelial detection, 136 (96.5%) were detected in at least 1% of malignant epithelial cells (Figure 4C; Table S6). Because detection estimates were unavailable for most candidates, single-cell expression was retained as supporting evidence rather than used as an exclusion criterion.

For transcript-resolved screening, we selected Caco-2 and HCT116, two CRC models with available transcriptome-wide long-read datasets and experimental tractability for downstream characterization. CCLE/DepMap expression analysis identified 488 disease-supported lncRNAs expressed at ≥1 TPM in Caco-2 and 402 in HCT116, with 577 detected in at least one model and 313 detected in both (Figure 4D; Table S6). After excluding *CRNDE*, which was examined separately in Figure 2, 576 lncRNAs entered systematic long-read analysis. Under the predefined processed-transcript criteria, 356 candidates had support from at least one qualifying processed long-read molecule containing an annotated splice junction, whereas 220 lacked qualifying processed-transcript evidence at the available sequencing depth (Figure 4E; Table S6).

We next determined whether the genomic regions overlapping published snoRNP-protein CLIP signals were retained within these processed RNA molecules. To distinguish gene-level transcript recovery from retention of discrete RNA regions, we reconstructed genomic CLIP-supported components and evaluated their retention within processed long-read alignments. Of the 356 candidates with processed-transcript support, 355 were evaluable for component-level analysis, encompassing 4,618 reconstructed CLIP-supported components (Figure 4F; Table S6). Applying the predefined retention criteria identified 190 components across 66 lncRNA loci, with retention of the same genomic component supported by processed long-read molecules in both Caco-2 and HCT116. Exclusion of two components at the SNHG17 locus because of unresolved coordinate projections yielded a final QC-filtered set of 188 retained components across 65 lncRNA loci. These components had a median length of 36 nt, substantially shorter than the 1,689-nt *CRNDE* UCE-containing region examined in Figure 2, reflecting the different sequence scales assessed by the two analyses. Length-sensitivity analyses of the QC-filtered set retained 161 components across 59 genes at ≥30 nt and 12 components across 10 genes at ≥70 nt (Figure S4C). Together, these analyses identified 65 CRC-associated lncRNA loci containing CLIP-supported regions retained within processed transcripts in both cellular models, extending the retained-region architecture exemplified by *CRNDE*^UCE^ to additional lncRNAs.

### 2.5. Representative CRC-associated lncRNAs exhibit distinct transcript architectures and nucleolar localization patterns

Having identified CRC-associated lncRNAs retaining snoRNP-protein CLIP-supported regions within processed transcripts, we next examined their transcript-processing architectures and subcellular distribution. Using splice-aware long-read alignments, we classified individual CLIP-supported genomic components according to their retention, excision, mixed processing, or unresolved status (Figures 5A; Table S7). As a reference for established intronic snoRNA-host processing, we examined *GAS5*, which harbors multiple annotated snoRNAs^8^. Among ten snoRNA-overlapping CLIP components, six exhibited predominantly excised architectures in Caco-2, with excision fractions ranging from 90.3% to 99.7% among reads assigned to either retained or excised states. The remaining four components showed lower excision fractions (1.0%-20.0%), although substantially fewer reads could be assigned to either resolved state (20-99 versus 572-581 for the excision-dominant components), limiting interpretation of their processing patterns. These observations demonstrated that the analysis could distinguish excised snoRNA-containing regions and highlighted the importance of resolving processing outcomes at the level of individual RNA components.

**Figure 5.**
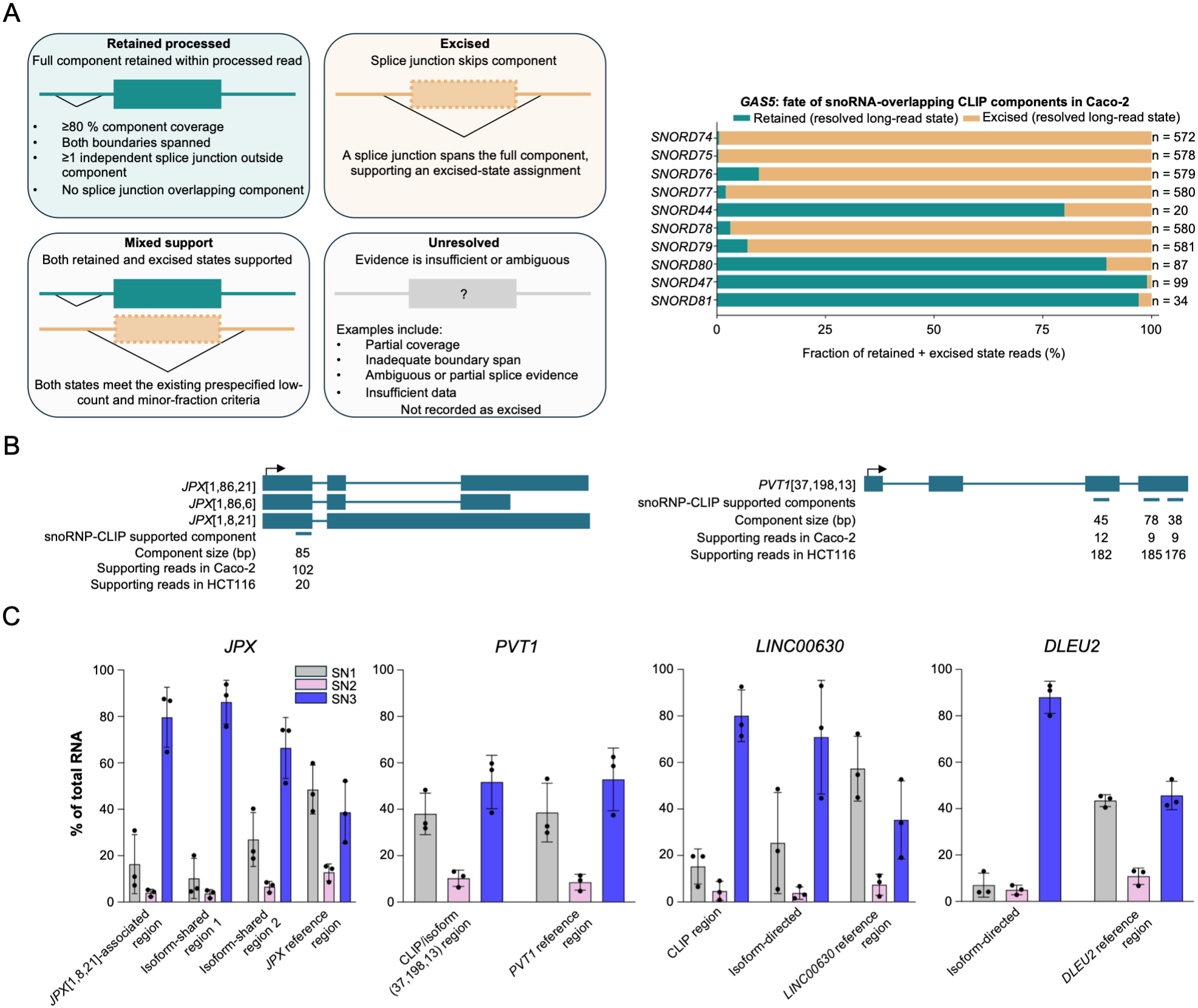
Representative CRC-associated lncRNAs exhibit distinct transcript architectures and nucleolar localization patterns. (A) Diagram showing classification of CLIP components. Retained processed components had ≥80% component coverage, both boundaries spanned, ≥1 independent splice junction outside component, and no splice junction overlapping the component. Excised components had a splice junction spanning the full component. Mixed support components had both retained and excised states supported, and unresolved components lacked sufficient evidence, and were not classified as excised. The fraction of *GAS5* CLIP components classified as retained or excised is shown as an illustrative example. (B) Transcript diagrams of *JPX* and *PVT1*, two lncRNAs selected for isoform-directed characterization. Transcript architectures supported by long-read data and snoRNP-CLIP supported components are shown. (C) Region-directed RT-qPCR analysis of *JPX*, *PVT1*, *LINC00630*, and *DLEU2* RNA distributions across pre-ribosome sequential extraction fractions in Caco-2 cells. *JPX* assays targeted a *JPX*[1,8,21]-associated region, two regions shared among selected *JPX* transcript models, and a broader *JPX* reference region. Reference-region assays for each locus served as broader transcript-region comparators and were not assumed to detect all transcript isoforms. None of the JPX amplicons directly overlapped the 85-nt CLIP-supported component shown in Figure 5B. These assays measure RNA populations containing the targeted regions rather than uniquely identifying complete transcript isoforms. Data are presented as mean ± SD (n=3 biological replicates).

To further characterize selected CRC-associated lncRNAs, we prioritized *JPX*, *PVT1*, *LINC00630*, and *DLEU2* based on their processed-transcript evidence, CRC disease association, and suitability for region-directed RT-qPCR analysis (Table S7). Among the 65 QC-filtered lncRNAs exhibiting stringent dual-model component retention, *JPX* and *PVT1* were prioritized for detailed transcript-architecture characterization because both retained identical snoRNP-protein CLIP-supported regions within processed transcripts from Caco-2 and HCT116, with shared complete splice-junction chains (Figure 5B). *LINC00630* combined tumor-upregulation and clinical-association evidence with membership in a ribosome-and RNP-related WGCNA module and stringent retention of five CLIP-supported components across both models, although its lower expression and lack of a shared complete splice-junction chain limited isoform-level resolution. *DLEU2*, which also mapped to a ribosome-related WGCNA module, was included as an expressed candidate with retention evidence in a distinct transcript-annotation context. Further isoform-level analysis identified an 85-nt CLIP-supported component in *JPX*, retained in 102 Caco-2 and 20 HCT116 processed reads, with five shared splice-junction chains, three of which matched ENCODE4 transcript annotations^24^. In *PVT1*, three CLIP-supported components of 45, 78, and 38 nt were retained across both models, with 9-12 supporting reads per component in Caco-2 and 176-185 in HCT116. Each *PVT1* component was associated with a shared splice-junction chain, although none of these shared chains exactly matched a complete ENCODE4 transcript annotation. These candidates provided an opportunity to examine the subcellular distribution of selected RNA regions relative to broader transcript populations from the same loci.

To determine whether selected RNA populations derived from the same lncRNA locus exhibited distinct subcellular distributions, we examined *JPX*, *PVT1*, *LINC00630*, and *DLEU2* using the PSE approach described above. Given that C/D snoRNP core proteins associate with guide snoRNAs and contribute to their nucleolar localization, where these complexes function in pre-rRNA maturation^7,25^, we reasoned that nucleolar compartmentalization would provide a relevant cellular context for characterizing these candidate RNA populations. Region-directed RT-qPCR assays were used to examine selected transcript-associated or CLIP-annotated regions, as applicable, alongside broader transcript-reference regions (Figure 5C).

In Caco-2 cells, *JPX* showed a striking distinction between region-directed and reference-region measurements. Whereas the broader *JPX* reference-region signal was distributed between SN1 and SN3, all three selected region-directed signals showed preferential enrichment in SN3 (Figure 5C). This pattern was also observed in HCT116 (Figure S5A). Two of these assays targeted regions shared among the selected processed *JPX* transcript models, whereas the third targeted a region associated with *JPX*[1,8,21]. None of the three *JPX* amplicons directly overlapped the 85-nt CLIP-supported component identified by long-read analysis (Figure 5B; Table S8).

Selected region-directed RNA populations from *DLEU2* and *LINC00630* also showed greater SN3 representation than their corresponding broader transcript measurements in both Caco-2 and HCT116, although the extent of enrichment varied among assays (Figures 5C and S5A). In contrast, the selected *PVT1* region-directed assay showed a distribution comparable to the broader *PVT1* measurement in Caco-2, whereas greater SN3 representation of the targeted RNA population was observed in HCT116 (Figure S5A). Together, these findings reveal preferential core nucleolar enrichment of selected RNA populations from CRC-associated lncRNA loci that is not apparent from broader transcript measurements. These observations highlight the value of region-directed analysis for resolving differences in subcellular distribution among RNA populations derived from the same locus.

## 3. Discussion

Although numerous lncRNAs have been implicated in CRC, the transcript architectures and subcellular distributions of the RNA populations underlying these associations remain poorly characterized. Building on *CRNDE*^UCE^ as a functional precedent, we investigated the UCE-containing region in CRC and integrated CRC disease evidence, published C/D snoRNP-protein CLIP profiles from HEK293 cells, and long-read RNA sequencing to identify additional lncRNAs retaining CLIP-supported regions within processed transcripts. This approach identified 1,714 disease-supported lncRNAs overlapping published snoRNP-protein CLIP signals, including 65 loci with 188 CLIP-supported regions retained within processed transcripts in both CRC cellular models examined. Region-directed subcellular analysis further revealed preferential core nucleolar enrichment of selected RNA populations from *JPX*, *DLEU2*, and *LINC00630* relative to broader transcript-region measurements from the same loci. These observations highlight the value of examining RNA regions, transcript-processing architectures, and subcellular distributions beyond conventional locus-level expression analyses.

The molecular function of *CRNDE*^UCE^ was previously established in RCC, where its ultraconserved snoRNA-like element promotes 60S ribosomal subunit maturation by facilitating eIF6 recruitment to maturing pre-60S subunits^11^. Although *CRNDE* was originally identified through its marked upregulation in colorectal neoplasia^12^, the representation and processing of its UCE-containing region in CRC had not been systematically characterized. We detected this region in nearly all assessable primary CRC tumors. Although absolute UCE-region coverage correlated with total *CRNDE* expression, its relative representation varied across tumors. We also identified processed *CRNDE* transcripts retaining the UCE-containing region in Caco-2 cells, with an additional qualifying molecule recovered from SW620 at limited sequencing depth. Region-directed analysis further revealed preferential core nucleolar enrichment of UCE-containing RNA in Caco-2 cells, whereas its distribution differed in HCT116. Despite recurrent *CRNDE* upregulation in CRC, total *CRNDE* expression showed limited association with disease progression or clinical outcome, and UCE-region signal did not define a significant, region-specific ribosome-biogenesis transcriptional program. These findings underscore that gene-level expression, regional RNA representation, processed transcript retention, and subcellular localization provide distinct levels of molecular information. This distinction may be particularly relevant in CRC, which exhibits considerable molecular and cellular heterogeneity, including distinct transcriptional subtypes and malignant epithelial states^26,27^. Accordingly, the potential biological relevance of *CRNDE*^UCE^ in CRC may depend on the representation and processing of particular RNA populations rather than total *CRNDE* abundance alone. Determining whether these RNA populations are preferentially represented in specific CRC states and contribute to ribosome maturation in those contexts represents an important direction for future investigation.

Our identification of additional lncRNAs retaining CLIP-supported regions within processed transcripts provides a broader perspective on the diversity of RNA-processing architectures involving regions with snoRNP-protein CLIP evidence. Canonical intronic snoRNAs are generally excised from their host transcripts, consistent with the excision-dominant architectures observed for several snoRNA-containing regions of *GAS5* in our analysis. Alternative processing architectures have also been described, including sno-lncRNAs flanked by snoRNA sequences and snoRNA-ended, polyadenylated lncRNAs^28,29^. Other examples, including *SLERT* and *LoNA*, demonstrate that snoRNA-related sequences can be incorporated into longer functional RNAs involved in the regulation of rRNA transcription or modification^9,10^. In this context, our long-read analysis identified discrete regions overlapping published snoRNP-protein CLIP signals that were retained within processed transcripts from additional CRC-associated lncRNA loci, including *JPX* and *PVT1*. These retained regions were considerably shorter than the *CRNDE* UCE-containing functional region and represent distinct molecular features rather than established functional equivalents of *CRNDE*^UCE^. Nevertheless, their retention within processed transcript architectures identifies specific RNA regions for further molecular investigation. This is particularly relevant because biological effects associated with lncRNA loci may arise from the RNA product itself, the process of transcription or splicing, or underlying regulatory DNA elements^30–32^. Moreover, the previously described roles of *JPX* in XIST regulation or *PVT1* in MYC-associated regulation^33,34^ do not necessarily encompass the molecular properties of every RNA population produced from these loci. Our findings therefore expand the molecular characterization of these lncRNAs beyond their previously studied activities and provide a basis for investigating whether retained RNA regions contribute to additional biological functions.

A particularly important finding of this study is that assays targeting different regions of the same lncRNA locus can reveal distinct subcellular distribution patterns. RNA localization is closely linked to lncRNA processing and function, and previous studies have documented substantial heterogeneity in lncRNA localization across transcripts and cellular contexts^35–38^. More recently, isoform-resolved analyses have demonstrated that gene-level measurements can obscure differences in the nuclear and cytoplasmic distribution of individual lncRNA isoforms^39^. Our findings extend this perspective to the core nucleolar compartment in CRC cells. In both Caco-2 and HCT116, three region-directed *JPX* assays showed preferential enrichment in the core nucleolar fraction (SN3) relative to the broader *JPX* reference-region assay. Two of these assays targeted regions shared among the selected processed *JPX* transcript models, whereas the third targeted a region associated with the *JPX*[1,8,21] transcript model. These localization assays did not directly interrogate the 85-nt CLIP-supported component identified by long-read analysis (Figure 5B); instead, they measured distinct transcript-associated regions (Figure 5C; Table S8). Selected *DLEU2* and *LINC00630* RNA populations similarly showed greater SN3 enrichment relative to their corresponding broader transcript measurements in both CRC models, whereas *PVT1* exhibited a more cell-line-dependent pattern. These observations reveal that the choice of RNA region being measured can substantially influence the apparent subcellular distribution of a lncRNA locus. The nucleolus is spatially organized into functionally distinct compartments that coordinate successive stages of pre-rRNA synthesis, processing, and ribonucleoprotein assembly^40,41^. Preferential recovery of selected RNA populations in a fraction enriched for early pre-ribosomal species therefore provides a relevant cellular context for investigating their potential involvement in nucleolar RNA metabolism. More broadly, our findings illustrate how region-directed measurements can reveal subcellular heterogeneity that is not readily resolved by broader transcript measurements, emphasizing the importance of RNA-region-resolved characterization in studies of lncRNA biology.

Several distinctions are important when considering the functional implications of these findings. We used published snoRNP-protein CLIP profiles generated in HEK293 cells to annotate candidate RNA regions, rather than to establish snoRNP association in CRC cells. Similarly, tumor-associated expression and clinical associations were defined at the gene level and cannot automatically be attributed to individual processed transcripts. Although a subset of the disease-supported candidates was associated with ribosome- and RNP-related transcriptional programs, neither these associations nor nucleolar enrichment demonstrates a role in ribosome biogenesis. Establishing such functions will require direct assessment of snoRNP association and pre-rRNA interactions in CRC cells, together with selective perturbation of retained RNA regions and transcript-specific rescue to determine their contributions to rRNA processing, ribosomal subunit maturation, and cancer cell growth. The previously established function of *CRNDE*^UCE^ provides a mechanistic precedent for these investigations without implying that the additional candidates share its mechanism. Together, our findings identify processed transcripts retaining published snoRNP-protein CLIP-supported regions across CRC-associated lncRNA loci and reveal preferential core nucleolar enrichment of selected RNA populations that is not apparent from broader transcript measurements. These results establish a transcript- and region-resolved foundation for investigating the potential contributions of CRC-associated lncRNAs to nucleolar biology and ribosome biogenesis.

## 4. Materials and Methods

### 4.1 Bulk TCGA data acquisition and preprocessing

Gene-level raw counts and clinical metadata for TCGA colon adenocarcinoma and rectal carcinoma were downloaded from the Genomic Data Commons (GDC) data portal (Release 45.0) with TCGAbiolinks and combined as colorectal cancer (CRC). STAR-aligned counts against the GDC GRCh38 reference were used throughout. Matched primary tumor and solid tissue normal samples were identified by TCGA patient barcode, yielding 50 CRC pairs, while tumor-only analysis used 633 primary tumor samples.

### 4.2 Tumor-normal differential expression

Differential expression was performed with DESeq2 on raw counts, retaining genes with ≥10 counts in ≥2 samples. Paired comparisons used the design ∼ patient + condition with normal tissue as reference. Wald *P* values were adjusted by the Benjamini-Hochberg procedure, and log2FC were shrunken with apeglm via lfcShrink.

### 4.3 Clinical progression and survival analyses

Clinical analyses used primary tumor samples with available expression and clinicopathologic annotation. AJCC pathological T, N, and M annotations were collapsed to major stage categories, treated as ordinal, and filtered to exclude Tx, Nx, and Mx. Spearman rank correlation on patient-level log2(TPM+1) values assessed stage progression association. Overall survival and progression-free interval were analyzed by univariate Cox proportional hazards regression on patient-level log2(TPM+1) values, with hazard ratios reported per one pooled standard deviation of patient-level log2(TPM+1) increase.

### 4.4 RNA sequencing read coverage analysis

TCGA CRC primary tumor and paired normal data were assessed with recount3. The *CRNDE*^UCE^ functional core was defined as chr16:54,920,635-54,922,323 (1,689 bp). To computationally estimate completely unspliced, nascent *CRNDE* transcription, a 5’ first-intron control region was derived based on the experimental *CRNDE* architecture. A consensus non-exonic first-intron interval was computationally intersected across four GENCODE v26 five-exon transcript models (ENST00000501177, ENST00000502066, ENST00000559432, ENST00000559598) that accurately map the functional core to intron 3. The resulting consensus interval was trimmed by 100 base pairs at each edge to minimize boundary artifacts, yielding a final locked nascent control window at chr16:54,925,314-54,928,393. Automated fail-closed audits verified that neither the functional core nor the nascent control locus overlapped any non-*CRNDE* annotated transcript spans in the GENCODE v26 database. Raw and scaled coverage metrics were extracted using the megadepth binary. Samples were considered assessable if *CRNDE* raw read coverage ≥ 1x.

### 4.5 Public long read RNA-seq datasets and preprocessing

Publicly available PacBio Iso-Seq data from seven primary colorectal cancer (CRC)-associated tissue specimens, comprising four tumor and three adjacent normal samples, were obtained from the European Nucleotide Archive (ENA; accession PRJEB68074). Iso-Seq reads were processed using the PacBio Iso-Seq workflow to recover full-length, non-concatemeric (FLNC) transcript molecules. For each sequencing run, CCS reads were subjected to primer identification and removal using Lima with the Iso-Seq-specific settings (--isoseq --peek-guess) and a defined 10x Genomics 3′ primer reference. Reads were subsequently processed using isoseq tag with the T-12U-16B library design to identify transcript-associated molecular tags. Full-length non-concatemeric reads were generated using isoseq refine with poly(A)-tail detection required (--require-polya). FLNC reads from sequencing runs corresponding to the same biological specimen were combined at the sample level and deduplicated using isoseq dedup to generate the final molecule-level datasets. Deduplicated transcript molecules were converted to FASTQ format using SAMtools and aligned to the human GRCh38 primary-assembly reference genome using minimap2 in splice-aware mode (-ax splice -uf -C5 -- secondary=no). Resulting alignments were coordinate-sorted and indexed using SAMtools. Alignments overlapping the *CRNDE* genomic region (chr16:54,910,000–54,940,000; GRCh38) were extracted for downstream transcript-architecture analysis. The resulting locus-specific alignments were evaluated at the individual-molecule level using a custom Python classification workflow, with transcript retention, splice-junction structure, and strand orientation as described previously.

### 4.6 Identification and classification of CRNDE UCE-retaining transcripts

To characterize *CRNDE* transcript architectures at the individual-molecule level, long-read alignments were examined within the *CRNDE* locus (chr16:54,910,000–54,940,000; GRCh38), focusing on the 1,689-nt UCE-containing region (chr16:54,920,635–54,922,323). Only primary, same-strand alignments with mapping quality ≥20 were retained, excluding unmapped, secondary, supplementary, and strand-conflicting reads. UCE retention was calculated as the proportion of reference-aligned nucleotides overlapping the UCE interval, with near-complete retention defined as ≥95%. Splice junctions were identified from CIGAR reference-skipping operations (N) of ≥20 nt, and independent junctions were required to lie outside the UCE-containing region but within the *CRNDE* screening interval. Processed *CRNDE* UCE carriers were defined as molecules retaining ≥95% of the UCE and containing at least one independent splice junction. Molecules lacking these criteria were classified according to UCE overlap and splice-junction structure, including UCE-retaining without independent splice support, UCE-excising, and other *CRNDE* architectures.

### 4.7. C/D snoRNP-protein CLIP-seq overlap analysis

Publicly available CLIP-seq datasets for the C/D snoRNP proteins NOP56, NOP58, and FBL^16^ were used to assess protein-associated regions within the *CRNDE* UCE. Published CLIP peak coordinates were converted from hg19 to GRCh38 using UCSC liftOver and intersected with the previously defined *CRNDE* UCE-containing region and C/D-like interval. Overlap lengths were calculated in nucleotides, and NOP58 peaks from independent libraries were evaluated separately to assess reproducibility. Protein-associated regions overlapping the C/D-like interval were distinguished from peaks located elsewhere within the broader UCE-containing region.

### 4.8. AlphaFold3 structural modeling of the CRNDE-UCE C/D snoRNP complex

AlphaFold3^17^ was used to evaluate the structural compatibility of the *CRNDE* UCE RNA with the core C/D snoRNP proteins SNU13, NOP56, NOP58, and FBL. The UCE RNA sequence and protein sequences were provided as inputs for multicomponent structure prediction. The resulting model was examined to assess the spatial arrangement of the RNA and protein components, particularly the positioning of SNU13 relative to the C-box-like region and the organization of NOP56, NOP58, and FBL around the C/D-like element. Predicted structures were interpreted as evidence of structural compatibility rather than experimental confirmation of native snoRNP assembly or direct protein–RNA interactions.

### 4.9. Genome-wide snoRNP-protein CLIP overlap analysis

To characterize snoRNP-protein CLIP overlap across the lncRNA transcriptome, GENCODE v50 lncRNA annotations (GRCh38) were used to construct gene-level exonic intervals by merging overlapping exons across annotated transcripts. Published CLIP-seq peak datasets for FBL, NOP56, and NOP58^16^ were intersected with these intervals using a custom Python workflow. Overlaps were restricted to matching genomic strands when CLIP peak strand annotations were available. A minimum exonic overlap of ≥1 nucleotide was used for the primary analysis, with a ≥10-nucleotide threshold applied as a sensitivity analysis. CLIP overlap was recorded separately for each library, including independent NOP58 replicates, and aggregated at the gene level to determine the number of overlapping core snoRNP proteins. These annotations were used to identify lncRNAs with CLIP overlap from at least one, at least two, or all three core proteins.

### 4.10. Degree preserving permutation analysis of snoRNP-protein CLIP convergence

To determine whether snoRNP-protein CLIP convergence exceeded expectations based on overall CLIP exposure, we implemented a degree-preserving permutation analysis using a binary matrix of 34,866 GENCODE v50 lncRNAs across five CLIP libraries (NOP58 replicates A and B, NOP56, FBL, and FBL-MNase). A bipartite switching algorithm randomized CLIP-library assignments while preserving each lncRNA’s total number of positive libraries and each library’s total number of positive lncRNAs. Following 50,000 successful burn-in swaps, 1,000 permutations were generated with 5,000 successful swaps between consecutive samples (random seed 20261006). NOP58 and FBL replicate signals were combined by logical union for protein-level convergence analysis. Observed counts of lncRNAs overlapping at least two or all three core proteins, as well as those supported by both NOP58 libraries, were compared with their corresponding null distributions. Empirical upper-tail *P* values were calculated as (1 + number of permutations with counts ≥ observed)/(1 + total permutations). The analysis was repeated using a ≥10-nt CLIP overlap threshold to assess sensitivity.

### 4.11. Replicate-consistent NOP58 enrichment and conditioned-null analysis

To assess enrichment of replicate-consistent NOP58 CLIP overlap among CRC disease-supported lncRNAs, the 1,714 candidates were compared with the remaining 7,733 CLIP-overlapping lncRNAs using Fisher’s exact test. NOP58 replicate consistency was defined as exonic overlap in both independent NOP58 libraries. To account for differences in overall CLIP exposure, a degree-preserving permutation analysis was performed using the five-library binary CLIP matrix while preserving each lncRNA’s total number of positive libraries and each library’s overall prevalence. Candidate membership was held fixed across 1,000 permutations, and the number of candidates overlapping both NOP58 libraries was recorded for each permutation. The observed count was compared with the null distribution to calculate the observed-to-expected ratio and empirical upper-tail *P* value using a one-sided test with a +1 correction.

### 4.12. Gene set enrichment analysis

GSEA was performed using clusterProfiler against GO-BP gene sets. Genes were ranked by moderated t-statistic. To assess ribosome biogenesis enrichment, the GO:0042254 term was used. Enrichment for matched intronic controls was compared against *CRNDE*^UCE^ enrichment.

### 4.13. CCLE acquisition

CCLE transcriptomic profiles were retrieved from EBI (E-MTAB-2770) as TPM values. Missing values were set to 0 TPM and expression was transformed as log2(TPM+1). Cell lines annotated “colon”, “rectal”, or “colorectal” with a default DepMap model entry were retained as the CRC cohort. Gene-wise median expression in TCGA primary CRC tumors was used as the matching variable for background standardization. Background lncRNAs were assigned to fine expression-abundance strata based on these values. The proportion of background lncRNAs expressed in 1 model, 10%, 25%, or 50% of the 81 CRC models was weighted by the corresponding proportion of candidates in that stratum.

### 4.14. Weighted gene co-expression network analysis

WGCNA^42^ was performed on TCGA Primary Tumor samples from CRC, STAD, and ESCA. Raw counts were processed with edgeR, normalized by trimmed mean of M-values, and converted to log2 counts per million. Protein-coding and noncoding genes with CPM > 1 in ≥20% of samples were retained. Genes and samples with excessive missingness or zero variance were removed with WGCNA quality-control functions. A pooled network was built from the integrated three-cohort matrix. Soft-thresholding power was evaluated with pickSoftThreshold and set to 10 based on scale-free topology fit while retaining connectivity. Modules were detected with blockwiseModules using power = 10, networkType = “unsigned”, TOMType = “unsigned”, corType = “pearson”, minModuleSize = 30, mergeCutHeight = 0.25, deepSplit = 2, maxBlockSize = 5000, pamStage = TRUE, and pamRespectsDendro = FALSE.

### 4.15. Single-cell-derived metacell analysis

Metacell profiles and metadata were obtained from the TabulaTIME pan-cancer single-cell atlas (accessed December 10, 2025). CRC samples were extracted using the atlas cancer-type field. Major cell-type annotations were taken from the curated atlas field. Profiles annotated as malignant defined the malignant epithelial compartment, and all other annotated profiles were assigned to non-malignant compartments.

### 4.16. Processed transcript recovery and component-level CLIP-region retention

576 non-*CRNDE* disease-supported, snoRNP-protein CLIP-overlapping RNAs with ≥1 TPM in either HCT116 or Caco-2 entered long-read analysis. Genes with ≥1 qualifying processed long-read molecule containing an annotated splice junction were retained as 356 candidates with processed-transcript support. 355 of these had evaluable architecture. These genes were intersected with CLIP analysis to yield 4,618 reconstructed CLIP-supported components. After manual QC, 188 strictly retained components across 65 lncRNAs were retained.

### 4.17. Human cell culture

HCT116 cells (male) and Caco-2 cells (male) were cultured in Dulbecco’s Modified Eagle Medium (DMEM) supplemented with 10% fetal bovine serum and 1% penicillin/streptomycin. Cell lines were confirmed to be free of mycoplasma contamination.

### 4.18. RNA isolation and region-directed RT-qPCR

RNA was extracted from equal cell equivalents of the SN1, SN2, and SN3 fractions obtained by PSE. Fraction volumes were sampled at a volumetric ratio of 5:3:4, and RNA was extracted using the Direct-zol RNA Mini Prep (Zymo Research) with on-column DNase digestion according to the manufacturer’s instructions. Region-directed RT-qPCR was performed to assess the distribution of selected RNA populations across the three fractions. Primer sequences are provided in Table S8. All experiments were performed with three independent biological replicates, and data are presented as mean ± SD.

### 4.19. Pre-ribosome sequential extraction (PSE)

A total of 2 x 10^6^ cells were pelleted and resuspended in 500 μL SN1 buffer (20 mM HEPES, pH 7.5, 130 mM KCl, 10 mM MgCl_2_, 0.05% Igepal CA-630) supplemented with 1x complete protease inhibitor cocktail (Roche) and 100 U SUPERase·In RNase Inhibitor (Thermo Fisher Scientific). Lysates were pipetted 20 times and centrifuged at 3,800 rpm for 3 minutes at 4°C to collect the SN1 supernatant. The pellet was washed with 500 μL of SN1 buffer and resuspended in 300 μL SN2 buffer (10 mM HEPES, pH 7.5, 10 mM NaCl, 5 mM MgCl_2_, 0.1% Igepal CA-630, and 0.5 mg/mL heparin) supplemented with 100 U SUPERase·In RNase Inhibitor and 10 μL Turbo DNase (Thermo Fisher Scientific). Following a 10-minute incubation at room temperature with pipetting every 5 minutes, the sample was centrifuged at 11,500 rpm for 10 minutes at 4°C to collect the SN2 supernatant. The remaining pellet was resuspended in 400 μL of SN3 buffer (10 mM HEPES, pH 7.5, 10 mM NaCl, 4 mM EDTA, 0.1% Igepal CA-630, 0.04% sodium deoxycholate, 4 mM imidazole, 0.1 mg/mL heparin, and 1 mM DTT) supplemented with 1x complete protease inhibitor cocktail (Roche) and 100 U SUPERase·In RNase Inhibitor. Following a 20-minute incubation on a shaker at room temperature, the sample was pipetted up and down, vortexed, and centrifuged at 11,500 rpm for 10 minutes at 4°C. The resulting supernatant was collected as the SN3 fraction.

### 4.20. Western Blot

4X NuPAGE LDS Sample Buffer (Thermo Fisher Scientific) was added to PSE fractions from Caco-2 and HCT116 cells incubated for 5 minutes at 95°C. Proteins were separated on 4-12% Bis-Tris NuPAGE gels (Thermo Fisher Scientific) and wet transferred onto nitrocellulose membranes (0.45 μm, Cytiva). Blocking was performed in TBST containing 5% BSA for 1 hour at room temperature. Membranes were incubated with NCL(Abcam, Cat#ab22758), α-tubulin (Sigma, Cat#158573), FBL (Proteintech, cat# 66985-I-Ig), and H3 (Proteintech, Cat#17168-1-AP) primary antibodies overnight at 4°C, followed by incubation with IR dye-labeled Multi-rAb CoraLite Plus 750 secondary antibodies (Proteintech, Cat#RGAR006, RGAM006) for 1 hour at RT. Blots were imaged using a LICOR Odyssey imager.

## Author Contributions

J.-S.L. conceived and supervised the study. B.F., P.L., E.P., and J.-S.L. designed experiments and interpreted the results. B.F. and P.L. performed bioinformatic analyses. E.P. and B.F. performed experimental studies. R.J.B. and C.D. assisted with data interpretation and manuscript editing. B.F., P.L., E.P., and J.-S.L. wrote the manuscript.

## Institutional Review Board Statement

This study involved secondary analyses of publicly available, de-identified human transcriptomic and clinical datasets and experiments using established human cell lines. No human participants were recruited, and no new patient specimens were collected. Institutional ethical review and approval were not required for this study.

## Informed Consent Statement

Not applicable.

## Data Availability Statement

Publicly available datasets analyzed in this study were obtained from The Cancer Genome Atlas (TCGA COAD and READ), recount3, the Cancer Cell Line Encyclopedia (CCLE), TabulaTIME, and published C/D snoRNP-protein CLIP and long-read RNA-sequencing datasets. Primary CRC PacBio Iso-Seq data are available from the European Nucleotide Archive under accession PRJEB68074. Data supporting the analyses are provided in the article and Supplementary Figures S1-S5 and Tables S1-S8. Additional processed data and analysis scripts are available from the corresponding author upon reasonable request.

## Acknowledgments

We thank Dr. Weihang Chai for generously providing the Caco-2 cell line. We acknowledge the patients and investigators who contributed to the publicly available datasets analyzed in this study. This study utilized data generated by the TCGA Research Network: https://www.cancer.gov/tcga.

