## Supplemental figures for "Transcript-resolved analysis identifies lncRNA carrier architectures and selective nucleolar enrichment in colorectal cancer"

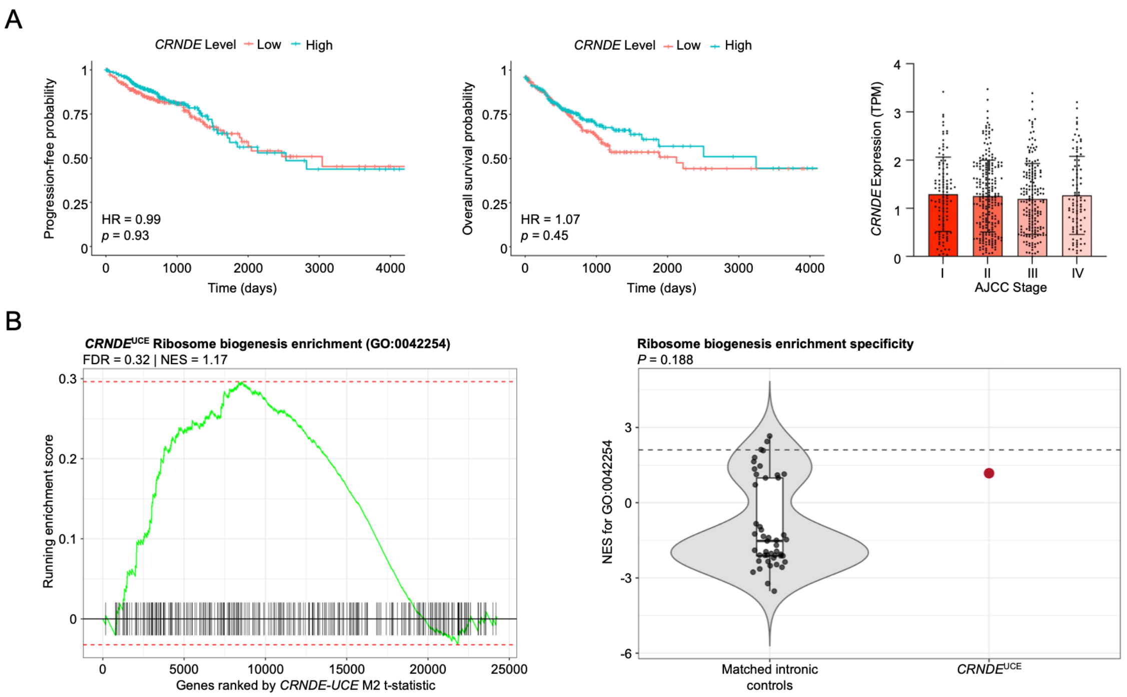


**Supplemental Figure S1. *CRNDE* is recurrently induced in colorectal cancer and the *CRNDE*$^{UCE}$ functional core is broadly detected in primary tumors**

(A) *CRNDE* clinical associations in primary CRC tumors. Left: Kaplan-Meier plot of progression-free interval stratified by median *CRNDE* expression. Center: Kaplan-Meier plot of overall survival using the same stratification. Right: *CRNDE* expression (TPM) across AJCC stages I-IV. (B) Association of the *CRNDE* UCE region with ribosome biogenesis in primary CRC tumors. Left: GSEA running-enrichment plot for ribosome biogenesis (GO:0042254; NES = 1.17; FDR = 0.32). Right: Comparison of the UCE-region enrichment score with matched intronic control regions (P = 0.188), showing no significant UCE-specific enrichment.

A

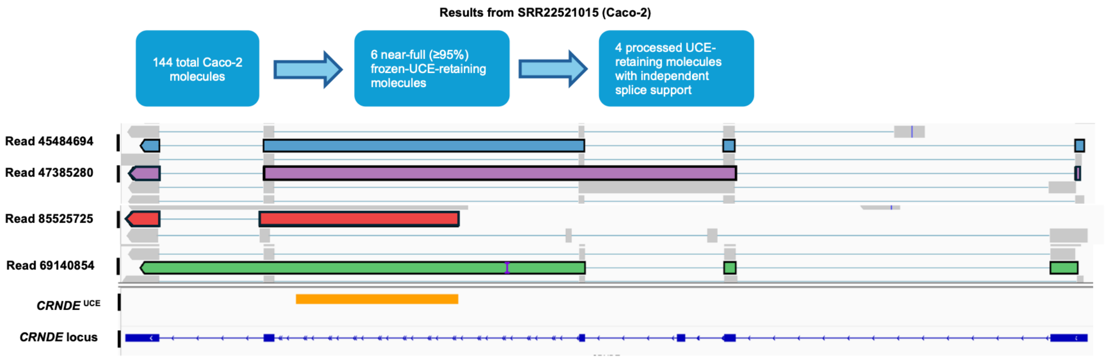


B

| Cell Line | High-confidence *CRNDE* reads | ≥ 95% UCE-retaining reads | Processed carriers | Carrier status |
|---|---|---|---|---|
| Caco-2 | 249 | 8 | 4 | Recovered |
| SW620 | 3 | 1 | 1 | Recovered (limited depth) |
| HCT116 | 32 | 0 | 0 | Not recovered |
| LoVo | 5 | 0 | 0 | Not recovered |
| SW480 | 2 | 0 | 0 | Not recovered |
| Colo320 | 0 | 0 | 0 | Unassessable |
| LS174T | 0 | 0 | 0 | Unassessable |

C

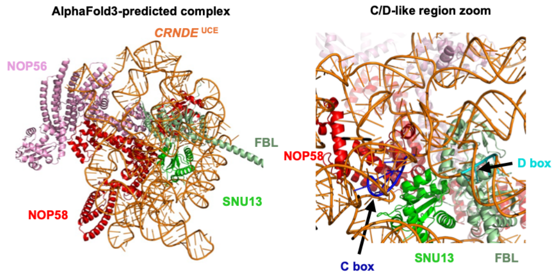


**Supplemental Figure S2. Long-read RNA-seq analysis identifies processed *CRNDE*$^{UCE}$ carriers in CRC with orthogonal support for C/D snoRNP association**

(A) Representative Caco-2 long-read RNA-seq dataset (SRR22521015), illustrating *CRNDE* transcript structures and splice-junction connectivity across the ultraconserved element (UCE)-containing interval. Among 144 *CRNDE* molecules in this dataset, six retained ≥95% of the UCE-containing region, and four had an independent splice junction and met the processed-carrier definition. The UCE interval is indicated for reference.

(B) Summary of processed *CRNDE* UCE-carrier recovery across CRC cell lines. In Caco-2, eight of 246 UCE-assessable *CRNDE* reads retained ≥95% of the UCE-containing region, including four with independent splice-junction support. SW620 yielded one qualifying carrier among three high-confidence *CRNDE* reads. $\textit{CRNDE}^{\mathrm{UCE}}$ carriers were not recovered in HCT116, LoVo, or SW480 and were not assessable in Colo320 and LS174T.

(C) AlphaFold3-predicted *CRNDE* UCE RNA–C/D snoRNP complex. Left: multicomponent structural model of *CRNDE* UCE RNA (orange), SNU13 (green), NOP56 (pink), NOP58 (red), and FBL (light green). Right: enlarged view of the C/D-like region, highlighting the predicted positioning of SNU13 near the C-box-like motif and the arrangement of the other proteins. The model suggests structural compatibility but does not establish native complex formation or experimentally validated RNA–protein interactions.

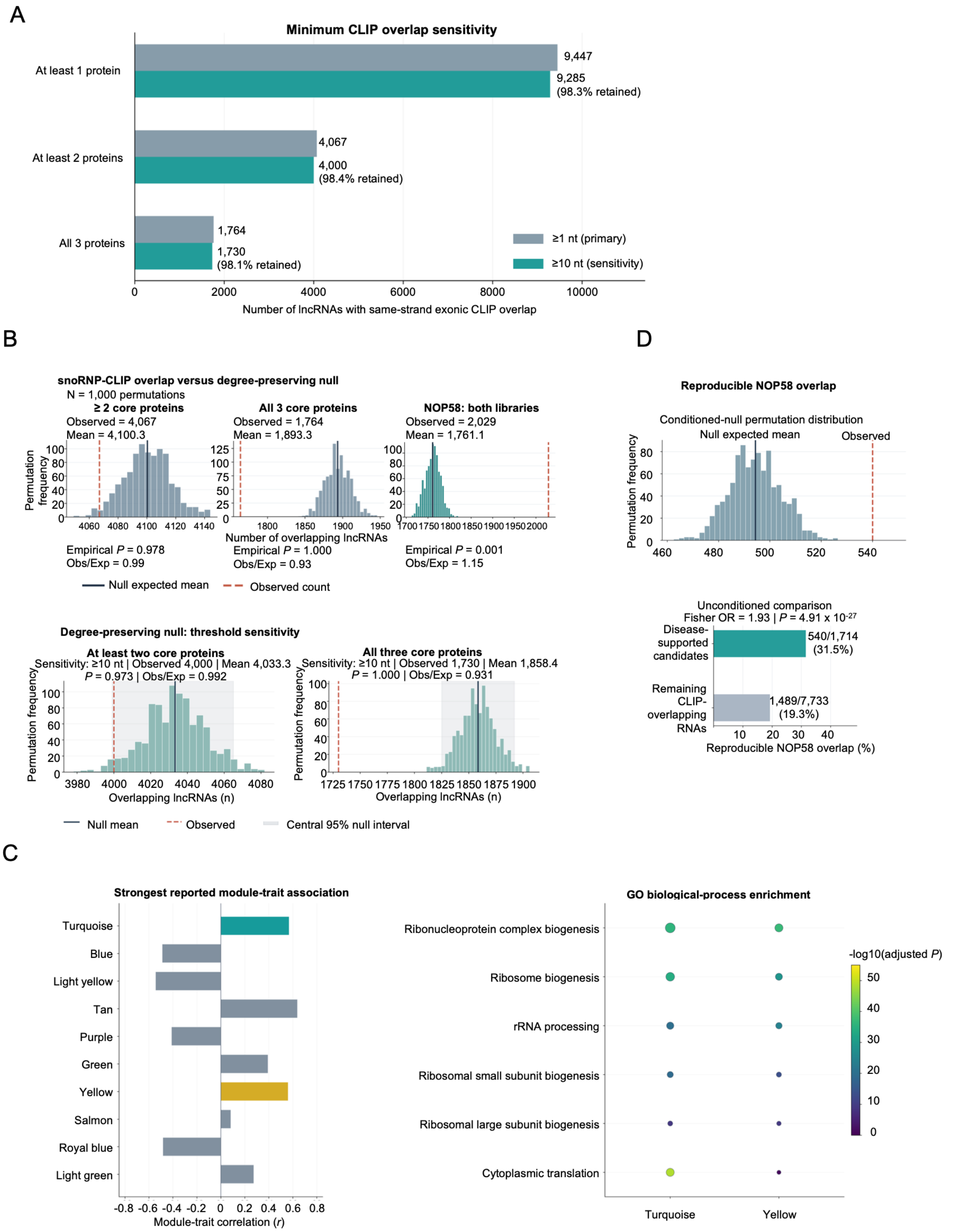


**Supplemental Figure S3. Integration of snoRNP-protein CLIP overlap with independent CRC disease evidence identifies candidate lncRNAs**

(A) Minimum CLIP-overlap sensitivity. Numbers of GENCODE v50 lncRNAs with same-strand exonic overlap of at least one, at least two, or all three core snoRNP proteins (FBL, NOP56 and NOP58), using ≥1 nt (primary analysis) or ≥10 nt (sensitivity analysis).

(B) Degree-preserving null analysis of snoRNP-protein CLIP overlap. Top: observed versus null-expected counts for ≥ 2 proteins (4,067 versus 4,100.3; empirical $P = 0.978$), all three proteins (1,764 versus 1,893.3; $P = 1.0$), and both NOP58 libraries (2,029 versus 1,761; $P = 0.001$). Bottom: sensitivity of the ≥ 2 proteins and all-three-proteins results to the ≥10-nt overlap threshold. The 1,000 bipartite permutations preserved lncRNA-level CLIP-library exposure and library-level prevalence; upper-tail empirical *P* values were calculated using a +1 correction.

(C) WGCNA context of CRC disease-supported lncRNAs. The strongest reported module-trait associations and GO biological-process enrichment provide complementary transcriptional-network context, including ribosome-, translation- and ribonucleoprotein-related processes.

(D) Replicate-consistent NOP58 overlap in CRC disease-support lncRNAs. Top: conditioned-null permutation distribution with 540 observed versus 494.4 expected candidates overlapping both NOP58 libraries. Bottom: proportions with replicate-consistent NOP58 overlap among disease-supported candidates (540/1,714; 31.5%) and the remaining CLIP-overlapping lncRNAs (1,489/7,733; 19.3%). The unconditioned comparison yielded Fisher's odds ratio of 1.93 ($P = 4.91 \times 10^{-27}$). The conditioned null accounts for overall CLIP-library exposure but does not explicitly control for transcript abundance.

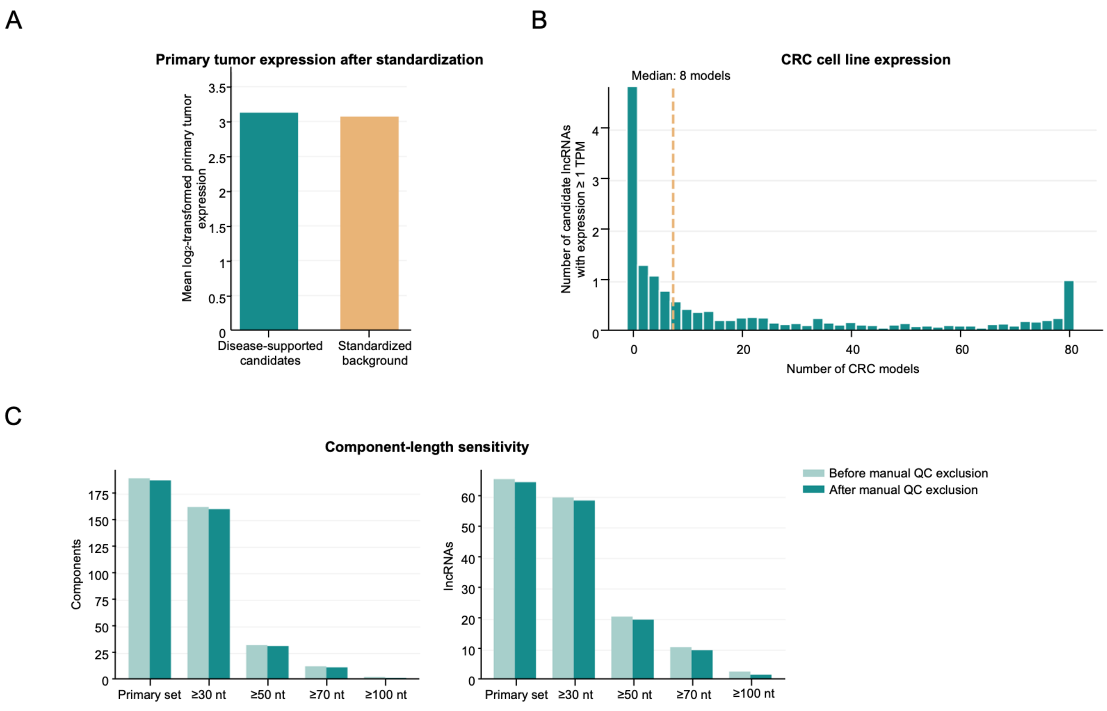


**Supplemental Figure S4. CRC cellular expression and long-read analysis identify transcript-resolved lncRNA carriers**

(A) Mean log2-transformed primary CRC tumor expression for disease-supported lncRNA candidates and the expression-standardized background lncRNA set.

(B) Distribution of the number of CRC cell-line models (of 81 examined) in which individual candidate lncRNAs were expressed at ≥ 1 TPM.

(C) Length-sensitivity analysis of retained CLIP-supported components and corresponding lncRNA loci, before and after manual quality-control (QC) exclusion, across the indicated component-length thresholds.

A

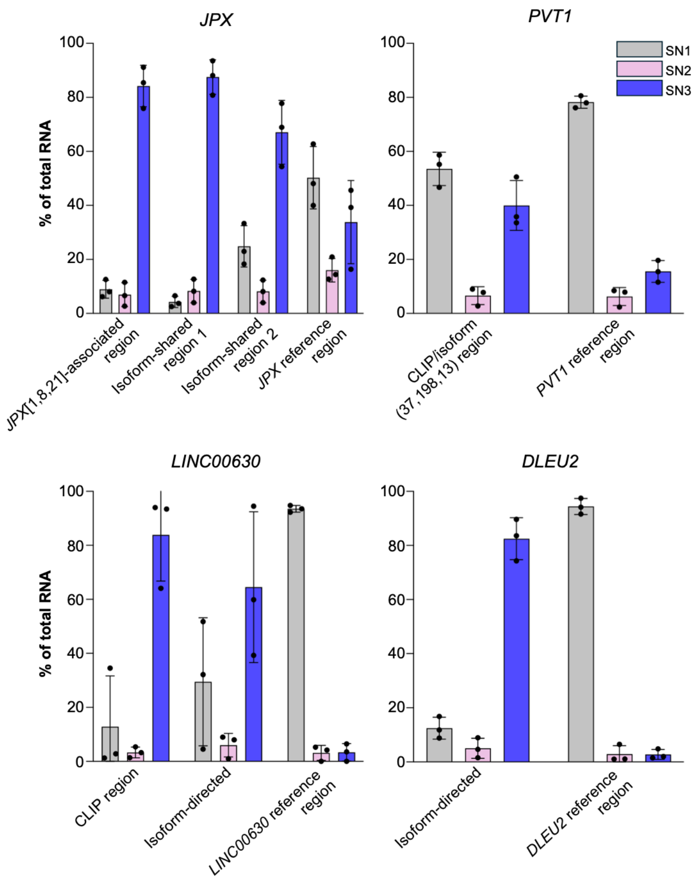


**Supplemental Figure S5. Representative CRC-associated lncRNAs exhibit region-dependent nucleolar localization patterns for HCT116 cells**

(A) Region-directed RT-qPCR analysis of *JPX*, *PVT1*, *LINC00630*, and *DLEU2* RNA distributions across pre-ribosome sequential extraction fractions in HCT116 cells. *JPX* assays targeted a *JPX*[1,8,21]-associated region, two regions shared among selected *JPX* transcript models, and a broader *JPX* reference region. A *PVT1* CLIP/isoform (37,198,13)-associated assay, *LINC00630* CLIP-region and isoform-directed assays, and a *DLEU2* isoform-directed assay were each compared with the corresponding reference-region assay. These comparators do not necessarily detect all transcript isoforms. Data are presented as mean ± SD (n=3 biological replicates).
